# MARK4 with an Alzheimer’s disease-related mutation promotes tau hyperphosphorylation directly and indirectly and exacerbates neurodegeneration

**DOI:** 10.1101/2020.05.20.107284

**Authors:** Toshiya Oba, Taro Saito, Akiko Asada, Sawako Shimizu, Koichi M. Iijima, Kanae Ando

## Abstract

Accumulation of the microtubule-associated protein tau is associated with Alzheimer’s disease (AD). In AD brain, tau is abnormally phosphorylated at many sites, and phosphorylation at Ser262 and Ser356 play critical roles in tau accumulation and toxicity. Microtubule-affinity regulating kinase 4 (MARK4) phosphorylates tau at those sites, and a double *de novo* mutation in the linker region of MARK4, ΔG316E317InsD, is associated with an elevated risk of AD. However, it remains unclear how this mutation affects phosphorylation, aggregation, and accumulation of tau and tau-induced neurodegeneration. Here, we report that MARK4^ΔG316E317D^ increases the abundance of highly phosphorylated, insoluble tau species and exacerbates neurodegeneration via Ser262/356-dependent and -independent mechanisms. Using transgenic *Drosophila* expressing human MARK4 (MARK4^wt^) or a mutant version of MARK4 (MARK4^ΔG316E317D^), we found that co-expression of MARK4^wt^ and MARK4^ΔG316E317D^ increased total tau levels and enhanced tau-induced neurodegeneration, and that MARK4^ΔG316E317D^ had more potent effects than MARK4^wt^. Interestingly, the *in vitro* kinase activities of MARK4^wt^ and MARK4^ΔG316E317D^ were similar. Blocking tau phosphorylation at Ser262 and Ser356 by alanine substitutions protected tau from the effects of MARK4^wt^, but not from MARK4^ΔG316E317D^. While both MARK4^wt^ and MARK4^ΔG316E317D^ increased the levels of oligomeric forms of tau, MARK4^ΔG316E317D^ further boosted the levels of tau phosphorylated at several sites other than Ser262/356 and increased the detergent insolubility of tau *in vivo*. Together, these findings suggest that MARK4^ΔG316E317D^ increases tau levels and exacerbates tau toxicity via an additional gain-of-function mechanism, and that modification in this region of MARK4 may impact disease pathogenesis.

## Introduction

Accumulation of the microtubule-associated protein tau is a pathological hallmark of AD (3–8). Tau accumulated in brains of AD patients is misfolded and highly phosphorylated by a number of kinases (5–7, 9). Among them, kinases that belong to the conserved Par-1/microtubule-affinity–regulating kinase (MARK) family phosphorylate tau within the microtubule-binding repeats at Ser262 and Ser356. Phosphorylation of tau at these sites regulates its microtubule binding, intracellular localization, and protein–protein interactions (10–16), and is also believed to play an initiating role in tau abnormality (13–19).

Mammals have four Par-1/MARK family members, MARK1–4 (3). MARK4 dysregulation has been proposed to play a role in the pathogenesis of AD. MARK4 expression is elevated in the brains of AD patients, and its activity colocalizes with early pathological changes (20). MARK4 can potentiate tau aggregation *in vitro* (21). A significant single‐nucleotide polymorphism (SNP) mapped to MARK4 in a regional Bayesian GWAS of AD (22). Importantly, a *de novo* double mutation in MARK4, ΔG316E317InsD, has been linked to elevated risk of early-onset AD (23). Phosphorylation at Ser262 is higher when tau is co-expressed with mutant MARK4 than when co-expressed with wild-type MARK4 (23), suggesting that this mutation increases the risk of AD by promoting production of abnormally phosphorylated tau. However, it is not fully understood how this mutation alters the effects of MARK4 on tau metabolism and toxicity.

In this study, we used a *Drosophila* model to compare the effects of MARK4 carrying the ΔG316E317InsD mutation (MARK4^ΔG316E317D^) on tau accumulation and toxicity with those of wild-type MARK4 (MARK4^wt^). The results revealed that co-expression of MARK4^ΔG316E317D^ increases the abundance of highly phosphorylated and insoluble tau species, resulting in enhanced accumulation and toxicity of tau, via a novel gain of function mechanism.

## Results

### MARK4^ΔG316E317D^ increases the levels of phosphorylated tau and promotes tau toxicity to a greater extent than MARK4^wt^

To assess the differences between the effects of MARK4^wt^ and MARK4^ΔG316E317D^ on metabolism and toxicity of tau *in vivo*, we established transgenic fly lines carrying human MARK4^wt^ and MARK4^ΔG316E317D^ under the control of a Gal4-responsive UAS sequence (Figure 1a). MARK4^wt^ or MARK4^ΔG316E317D^ was co-expressed with human tau in the retina using the pan-retinal GMR-Gal4 driver. Western blotting revealed that these flies expressed MARK4^wt^ and MARK4^ΔG316E317D^ at similar levels (Figure 1a). MARK4 phosphorylates tau at Ser262 and Ser356 (24), and we previously reported that tau phosphorylation at Ser262 by Par-1, a member of the PAR-1/MARK family, stabilizes tau and increases total tau levels (13, 14). Both MARK4^wt^ and MARK4^ΔG316E317D^ increased the levels of tau, as well as tau phosphorylated at Ser262 and Ser356; MARK4^ΔG316E317D^ had stronger effects on both (Figure 1b). The prominent increase in tau phosphorylation at Ser356 by MARK4^ΔG316E317D^ is noteworthy, since tau phosphorylation at Ser356 in this model reflects abnormally elevated PAR-1/MARK activity (14). Co-expression of MARK4^wt^ or MARK4^ΔG316E317D^ did not affect the mRNA levels of tau (Figure 1c), indicating that the increase in tau levels was not due to elevated transcription of the tau transgene.

**Figure 1.**
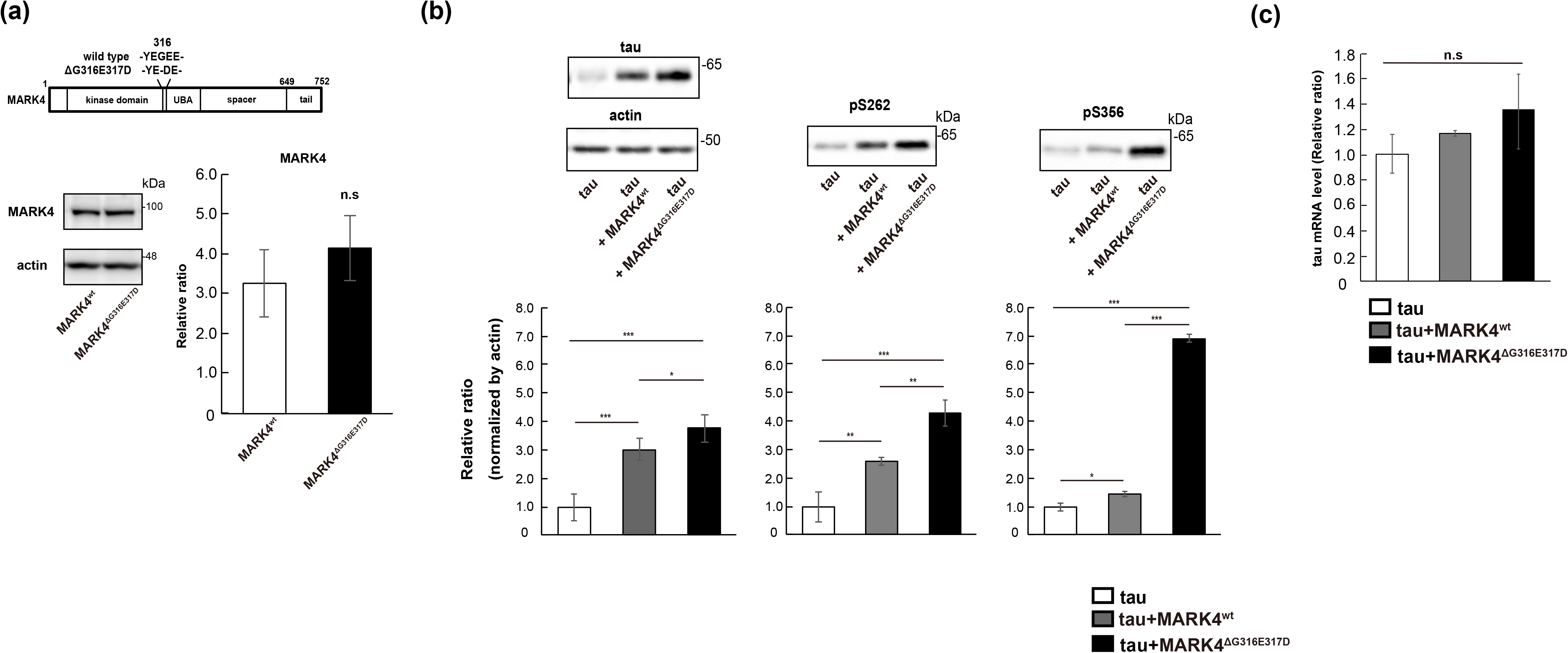
MARK4^ΔG316E317D^ increases the level of tau to a greater extent than MARK4^wt^. (a) (Top) Schematic representation of MARK4 and ΔG316E317D mutation. (Bottom) MARK4^wt^ and MARK4^ΔG316E317D^ are expressed at similar levels in fly eyes. MARK4 expression was driven by the pan-retinal driver GMR-Gal4. Levels of MARK4 in fly head lysates were examined by western blotting. Actin was used as a loading control. Representative blots (left panels) and quantification (right panels) are shown. (b) Co-expression of MARK4 increased the levels of phosphorylated tau and total tau, and co-expression of MARK4^ΔG316E317D^ increased the levels of total tau, pSer262-tau, and pSer356-tau in the eyes to a significantly greater extent than MARK4^wt^. Western blotting was performed with anti-tau antibody (total tau), anti-phospho Ser262 tau antibody (pSer262), or anti-phospho Ser356 antibody (pSer356). Representative blots (top) and quantification (bottom panels) are shown. (c) mRNA levels of tau expressed alone (tau), tau co-expressed with MARK4^wt^ (tau+ MARK4^wt^), and tau co-expressed with MARK4^ΔG316E317D^ (tau+ MARK4^ΔG316E317D^) were measured by quantitative PCR. Means ± SD; n = 4. n.s., p > 0.05; * p < 0.05; ** p < 0.01; *** p < 0.005 (one-way ANOVA and Tukey post-hoc test).

Next, we analyzed the effect of co-expression of MARK4 on tau toxicity. Expression of human tau in the eyes causes age-dependent and progressive neurodegeneration in the lamina, the first synaptic neuropil of the optic lobe containing photoreceptor axons (18). We found that flies co-expressing tau and MARK4^wt^ or MARK4^ΔG316E317D^ exhibited more neurodegeneration in the lamina than those expressing tau alone. Moreover, co-expression of MARK4^ΔG316E317D^ caused more prominent neurodegeneration than co-expression of MARK4^wt^ (Figure 2a and b). By contrast, expression of MARK4^wt^ or MARK4^ΔG316E317D^ alone, without tau, did not cause neurodegeneration at the same age (Figure 2a and b). Together, these results suggest that MARK4^ΔG316E317D^ promotes tau phosphorylation and accumulation and exacerbates tau toxicity to a greater extent than MARK4^wt^.

**Figure 2.**
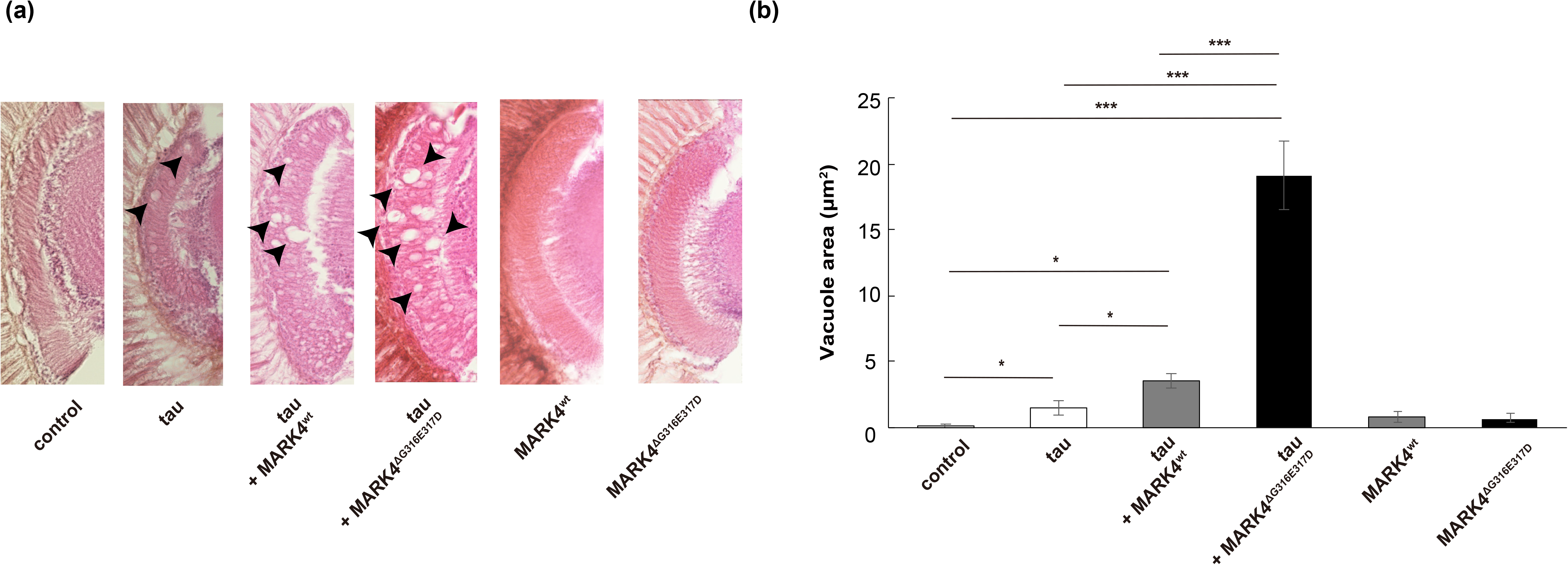
MARK4^ΔG316E317D^ increases tau toxicity to a greater extent than MARK4^wt^. (a) Co-expression of MARK4 increased tau-induced neurodegeneration, and MARK4^ΔG316E317D^ exerted a stronger effect than MARK4^wt^. Shown are lamina of flies carrying GMR-Gal4 driver alone (control), flies expressing tau (tau), flies co-expressing tau and MARK4^wt^ (tau+MARK4^wt^), or flies co-expressing tau and MARK4^ΔG316E317D^ (tau+MARK4^ΔG316E317D^). Expression of MARK4^wt^ (MARK4^wt^) or MARK4^ΔG316E317D^ (MARK4^ΔG316E317D^) alone did not cause neurodegeneration. Neurodegeneration is observed as vacuoles, indicated by arrows. (b) Quantification of vacuole area. Means ± SD; n = 5. n.s., p > 0.05; * p < 0.05; ** p < 0.01; *** p < 0.005 (one-way ANOVA and Tukey post-hoc test).

### MARK4^wt^ and MARK4^ΔG316E317D^ have similar kinase activities in vitro

A previous study reported that when HEK293 cells were co-transfected with tau and either MARK4^wt^ or MARK4^ΔG316E317D^, cells transfected with MARK4^ΔG316E317D^ exhibited a significant increase in tau Ser262 phosphorylation relative to those transfected with MARK4^wt^ (23). The authors interpreted this as evidence that the *de novo* mutation in MARK4 increases the ability of MARK4 to phosphorylate tau on Ser262 more efficiently (23). However, it has not yet been tested whether MARK4^ΔG316E317D^ has higher kinase activity than MARK4^wt^ To explore this possibility, we carried out an *in vitro* kinase assay. Specifically, we expressed MARK4^wt^ or MARK4^ΔG316E317D^ in HEK293 cells, immunoprecipitated MARK4 proteins from cell lysates, and measured their kinase activities using recombinant tau as a substrate. MARK4^wt^ or MARK4^ΔG316E317D^ had similar kinase activities (Figure 3), indicating that the higher level of tau accumulation in cells expressing MARK4^ΔG316E317D^ was not due to a difference in kinase activity.

**Figure 3.**
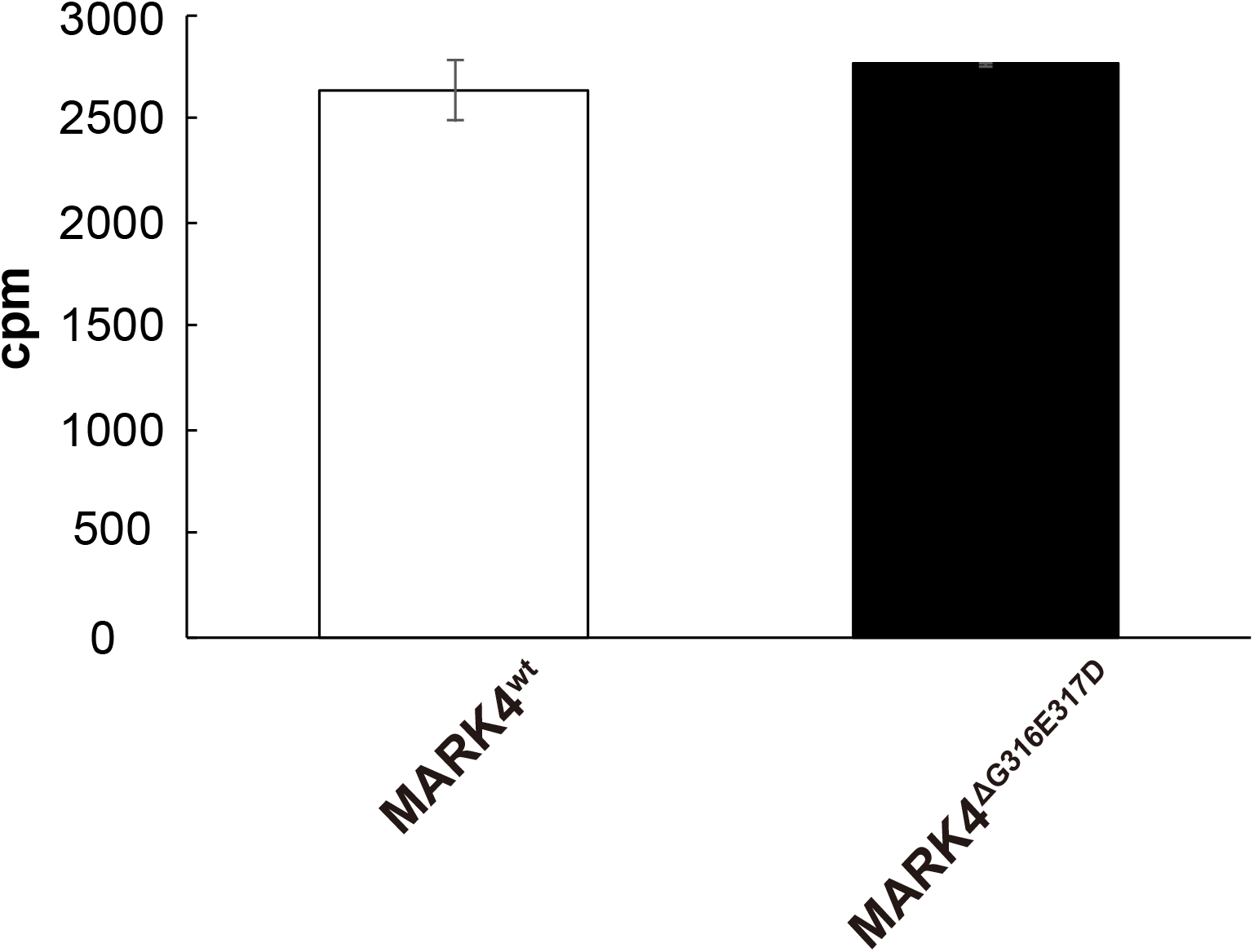
MARK4^wt^ or MARK4^ΔG316E317D^ have similar kinase activity *in vitro.* MARK4^wt^ and MARK4^ΔG316E317D^ expressed in HEK293 cells were immunoprecipitated and subjected to *in vitro* kinase assays. Incorporation of ^32^P in recombinant tau protein is expressed as means ± SEM (n = 3. n.s., p > 0.05).

### MARK4^ΔG316E317D^, but not MARK4^wt^, increases tau levels and exacerbates tau toxicity in a Ser262/356-independent manner

We previously reported that Par-1 overexpression causes tau accumulation via its phosphorylation at Ser262 and Ser356, thus enhance neurodegeneration (13, 14). To determine whether the exacerbation of tau accumulation and toxicity by MARK4^wt^ or MARK4^ΔG316E317D^ is also mediated by tau phosphorylation at Ser262 and Ser356, we used a tau mutant in which both of those sites are replaced by alanines (S2A) (25). Similar to Par-1 (13, 14), expression of MARK4^wt^ did not increase the level of S2A tau (Figure 4a). In addition, MARK4^wt^ did not exacerbate neurodegeneration caused by S2A tau (Figure 4b), suggesting that MARK4^wt^ increases tau levels and tau toxicity through tau phosphorylation at Ser262 and Ser356. By contrast, MARK4^ΔG316E317D^ significantly increased the levels of S2A tau (Figure 4a), and co-expression of MARK4^ΔG316E317D^ with S2A tau exacerbated neurodegeneration (Figure 4b). These results suggest that co-expression of MARK4^ΔG316E317D^ increases tau levels and tau toxicity via an additional gain-of-function mechanism that does not involve phosphorylation at Ser262 and Ser356.

**Figure 4.**
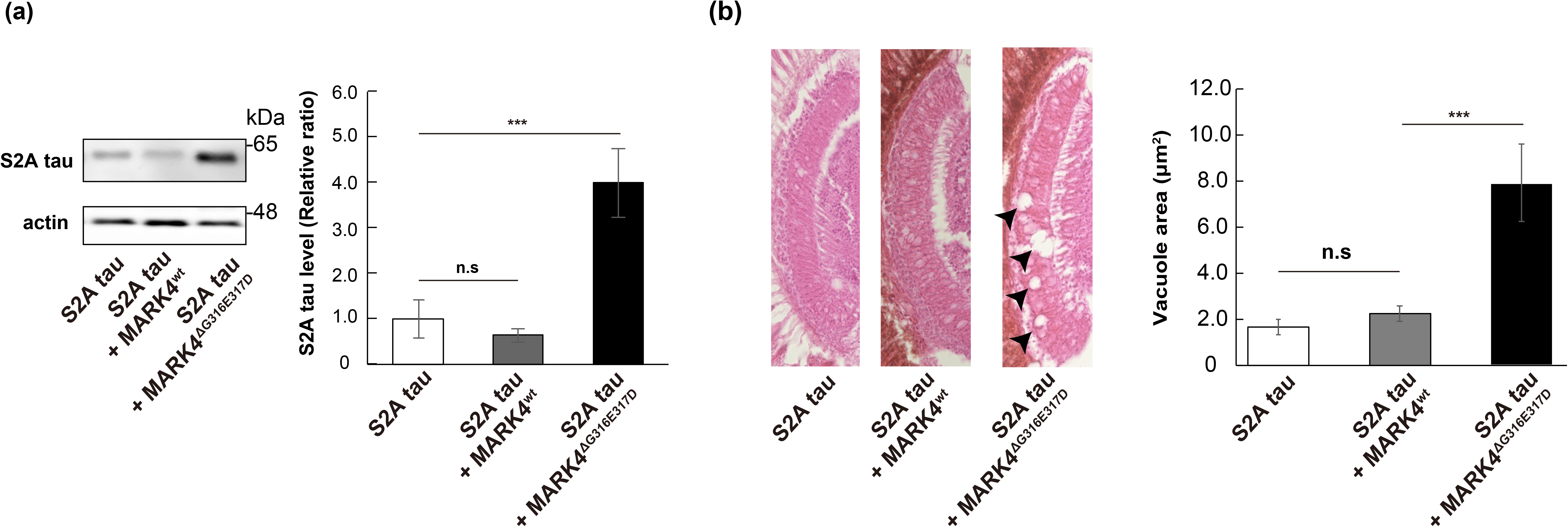
MARK4^ΔG316E317D^ increases tau levels and tau toxicity in a Ser262/356-independent manner. (a) MARK4^ΔG316E317D^ positively regulated tau levels in a Ser262/356-independent manner. Western blots of fly heads expressing S2A tau alone or co-expressing S2A tau with either MARK4^wt^ (S2A tau+MARK4^wt^) or MARK4^ΔG316E317D^ (S2A tau+MARK4^ΔG316E317D^). (b) Co-expression of S2A tau with MARK4^ΔG316E317D^, but not with MARK4^wt^, enhanced neurodegeneration. Shown are lamina of flies harboring expressing S2A tau (S2A tau), co-expressing S2A tau and MARK4^wt^ (S2A tau+MARK4^wt^), or co-expressing S2A tau and MARK4^ΔG316E317D^ (S2A tau+MARK4^ΔG316E317D^). Means ± SE; n = 5. n.s., p > 0.05; *** p < 0.005 (one-way ANOVA and Tukey post-hoc test).

### Expression of MARK4^wt^ or MARK4^ΔG316E317D^ does not disrupt overall protein degradation

To investigate the mechanisms underlying accumulation of tau caused by MARK4^ΔG316E317D^, we first sought to determine whether this effect is specific to tau protein. When co-expressed with firefly luciferase, MARK4^ΔG316E317D^ did not increase luciferase levels, indicating that the mutant protein does not cause non-specific accumulation of exogenously expressed proteins (Figure 5a).

**Figure 5.**
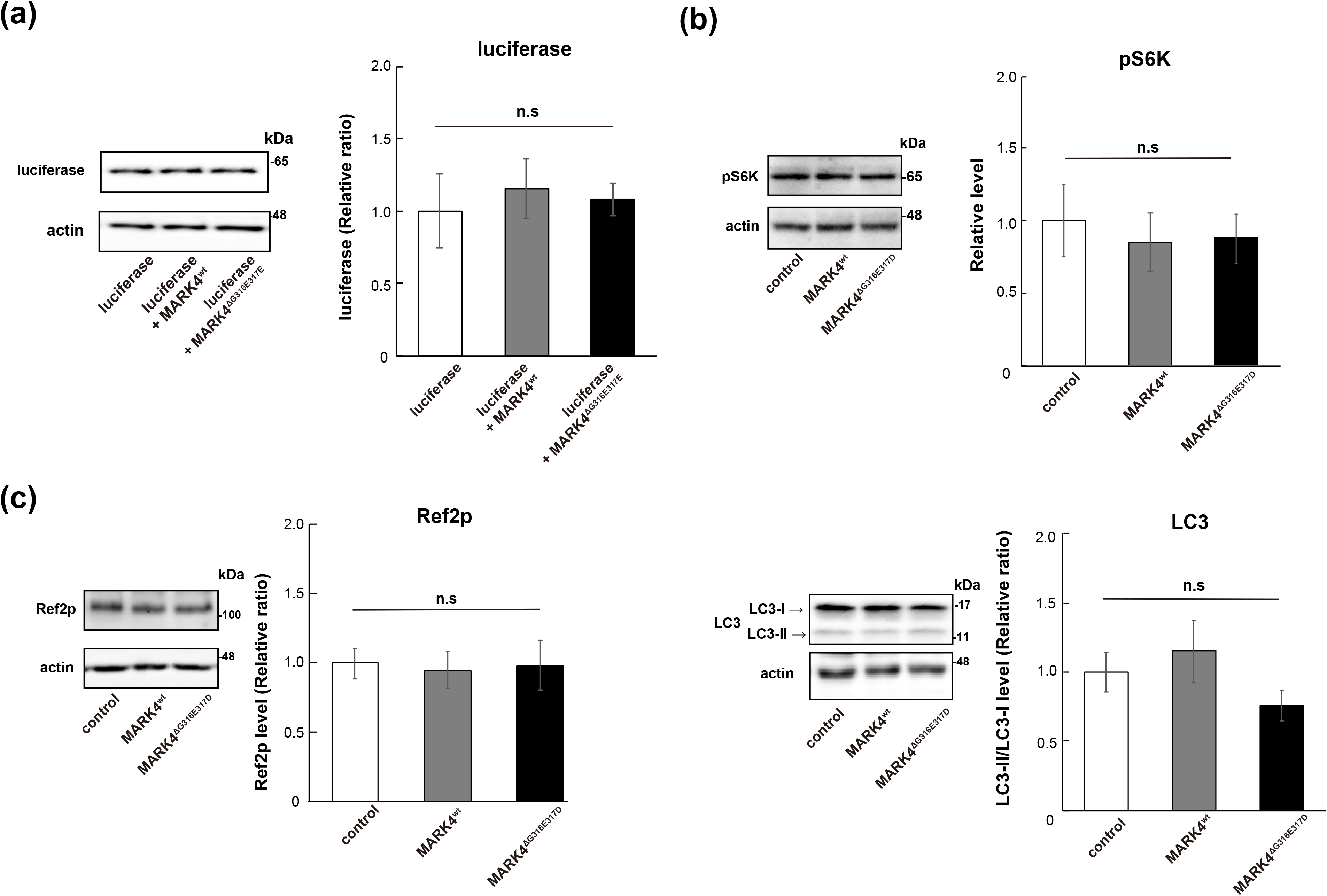
Neither MARK4^wt^ nor MARK4^ΔG316E317D^ inhibits the TOR pathway or increases autophagic activity. (a) MARK4^ΔG316E317D^ did not increase the levels of luciferase co-expressed in fly eyes. Western blots were performed on fly heads expressing luciferase alone (luciferase), co-expressing luciferase and MARK4^wt^ (luciferase+MARK4^wt^), or co-expressing luciferase and MARK4^ΔG316E317D^ (luciferase+MARK4^ΔG316E317D^). Actin was used as a loading control. Representative blots and quantitation are shown. Means ± SD; n = 4. n.s., p > 0.05; * p < 0.05; ** p < 0.01; *** p < 0.005 (one-way ANOVA and Tukey post-hoc test). (b-d) Expression of MARK4^wt^ or MARK4^ΔG316E317D^ did not promote autophagy. Western blots were performed on fly heads expressing control, MARK4^wt^, or MARK4^ΔG316E317D^. Blots were performed with anti-pS6K (pS6K) (b), anti-Ref2P (Ref2P) (c), and LC3 antibody (LC3) (d). Representative blots and quantitation are shown. Means ± SEM; n = 3. n.s., p > 0.05; * p < 0.05, *** p < 0.005 (one-way ANOVA and Tukey post-hoc test). Actin was used as a loading control.

We next examined whether MARK4^ΔG316E317D^ inhibits autophagic activity. MARK4 is known to negatively regulate the mechanistic target of rapamycin complex 1 (mTORC1) (26). mTORC1 is a conserved regulator of autophagy, which mediates degradation of a variety of proteins, including tau (27). To test whether MARK4^wt^ and MARK4^ΔG316E317D^ had distinct effects on mTOR signaling, we first investigated whether expression of MARK4^wt^ or MARK4^ΔG316E317D^ in the fly retina inhibits TOR signaling. We found that neither MARK4^wt^ nor MARK4^ΔG316E317D^ affected phosphorylation of S6K, a target of TOR (Figure 5b). In addition, neither MARK4^wt^ nor MARK4^ΔG316E317D^ affected LC3-II/LC3-I ratio or the levels of the autophagic substrate Ref2P (Figure 5c), suggesting that autophagic activity was not dampened. These results suggest that MARK4^ΔG316E317D^ causes accumulation of tau via a mechanism other than overall suppression of protein degradation.

### MARK4^ΔG316E317D^ increases the levels of highly phosphorylated tau species

Because hyperphosphorylation of tau is correlated with severity of tau pathology, we asked whether MARK4^ΔG316E317D^ increases the levels of phosphorylated tau to a greater extent than MARK4^wt^ *in vivo*. For this purpose, we used Phos-tag analysis for comprehensive quantitative profiling of the phosphorylation status of tau (28). *Drosophila* expresses a single member of the Par-1/MARK family, Par-1; we previously reported that Par-1 phosphorylates human tau proteins at Ser262 and Ser356, stabilizing them and promoting their subsequent phosphorylation at other sites (15, 18). To highlight the difference between the effects of MARK4^wt^ and MARK4^ΔG316E317D^ on tau phosphorylation in this model, we carried out this experiment in a Par-1–knockdown background. Tau protein expressed in the fly retina separated into several bands, indicating that it was phosphorylated in multiple patterns (Figure 6a). Interestingly, the phosphorylation forms of tau whose abundances increased differed depending on whether MARK4^wt^ or MARK4^ΔG316E317D^ was co-expressed. MARK4^wt^ increased the intensity of a faster-migrating band (tau_lower_), whereas co-expression MARK4^ΔG316E317D^ shifted the major bands higher (tau^upper^), indicating that the mutant protein increased the levels of highly phosphorylated forms of tau *in vivo* (Figure 6a).

**Figure 6.**
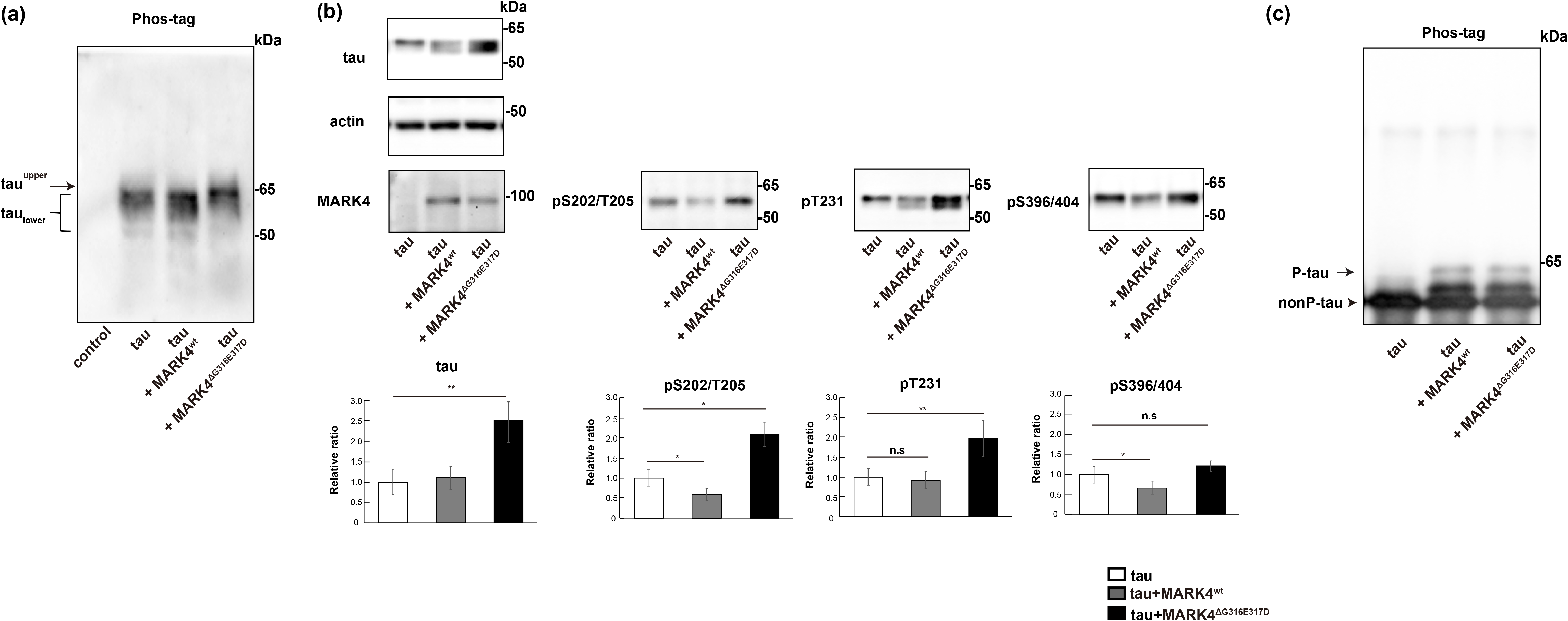
Co-expression of MARK4^ΔG316E317D^ increases tau phosphorylated at sites other than Ser262/356 *in vivo*. (a) Phosphorylation profile of tau expressed alone or co-expressed with MARK4^wt^ or MARK4^ΔG316E317D^ *in vivo*. Fly head extracts were separated by Phos-tag SDS-PAGE, followed by western blotting with anti-tau antibody. Co-expression of MARK4^wt^ increased the abundance of faster-migrating tau (tau_lower_). By contrast, co-expression of MARK4^ΔG316E317D^ increased the abundance of slower-migrating tau (tau^upper^). (b) Western blotting was performed using phospho-specific antibodies for SP/TP sites such as AT8 (pS202/205), anti-pThr231 (pT231), and PHF1 (pS396/404), as well as pan-tau antibody (total tau). Actin was used as a loading control. Representative blots (top) and quantification (bottom panels, normalized either by actin) are shown. Means ± SEM; n = 4. n.s., p > 0.05; * p < 0.05; ** p < 0.01; *** p < 0.005 (one-way ANOVA and Tukey post-hoc test). (c) MARK4^wt^ and MARK4^ΔG316E317D^ phosphorylated tau in the same pattern *in vitro*. MARK4^wt^ and MARK4^ΔG316E317D^ expressed in HEK293 cells were immunoprecipitated and incubated with recombinant tau. Tau proteins were separated by Phos-tag SDS-PAGE, followed by western blotting with anti-tau antibody.

We previously reported that the difference in migration speed among these bands is related to their phosphorylation levels: tau in the slower-migrating band (tau^upper^) is more highly phosphorylated than tau in the faster-migrating band (tau_lower_) (13). Tau has many serine and threonine sites followed by prolines, which are phopshorylated by proline-directed Ser/Thr kinases (SP/TP kinases) such as GSK3β, MAPK and cyclin-dependent kinase 5 (Cdk5) (29–32). Particularly, tau^upper^ is phosphorylated at SP/TP sites (13) that are not direct substrates of non-SP/TP kinases including Par-1/MARK (15, 33). To determine whether MARK4^ΔG316E317D^ promotes accumulation of tau phosphorylated at SP/TP sites to a greater extent than MARK4^wt^, we carried out western blotting using phospho-specific antibodies against tau phosphorylated at the following SP/TP sites: AT8 (Ser202, Thr205), Thr231, and PHF1 (Ser396, Ser404). Co-expression of tau and MARK4^wt^ did not increase, or rather slightly decreased, the levels of tau phosphorylated at these sites (Figure 6b). By contrast, co-expression of tau and MARK4^ΔG316E317D^ significantly increased the levels of tau phosphorylated at AT8 and Thr231 sites relative to expression of tau alone or co-expression of tau and MARK4^wt^ (Figure 6b). These results suggest that MARK4^ΔG316E317D^ promotes the accumulation of tau species phosphorylated at additional sites other than Ser262 and Ser356.

Although these SP/TP sites are not expected to be direct substrates of MARK4 (15, 33), we wondered whether the ΔG316E317D mutation might alter the kinases’ substrate specificity. To determine whether MARK4^ΔG316E317D^ directly phosphorylates those sites, we carried out an *in vitro* assay. We expressed MARK4^wt^ or MARK4^ΔG316E317D^ in HEK293 cells, immunoprecipitated the cell lysates, and incubated the immunoprecipitated proteins with recombinant tau; we then analyzed the phosphorylation pattern of tau using the Phos-tag method. Tau exhibited the same phosphorylation pattern whether it was incubated with MARK4^wt^ or MARK4^ΔG316E317D^ (Figure 6c), suggesting that MARK4^ΔG316E317D^ increases accumulation of tau species phosphorylated at SP/TP sites via an indirect mechanism *in vivo*.

### MARK4^wt^ and MARK4^ΔG316E317D^ both increase tau dimers and oligomers, and MARK4^ΔG316E317D^ also promotes accumulation of insoluble forms of tau

Tau can aggregate into higher-order structures that are prone to accumulate. Hence, we asked whether co-expression of MARK4^wt^ or MARK4^ΔG316E317D^ would affect the formation of small aggregates such as dimer and oligomers. Tau proteins in *Drosophila* retina exist mostly in the monomeric form (19, 34). We found that both MARK4^wt^ and MARK4^ΔG316E317D^ increased the levels of dimers and oligomers to similar extents (Figure 7a).

**Figure 7.**
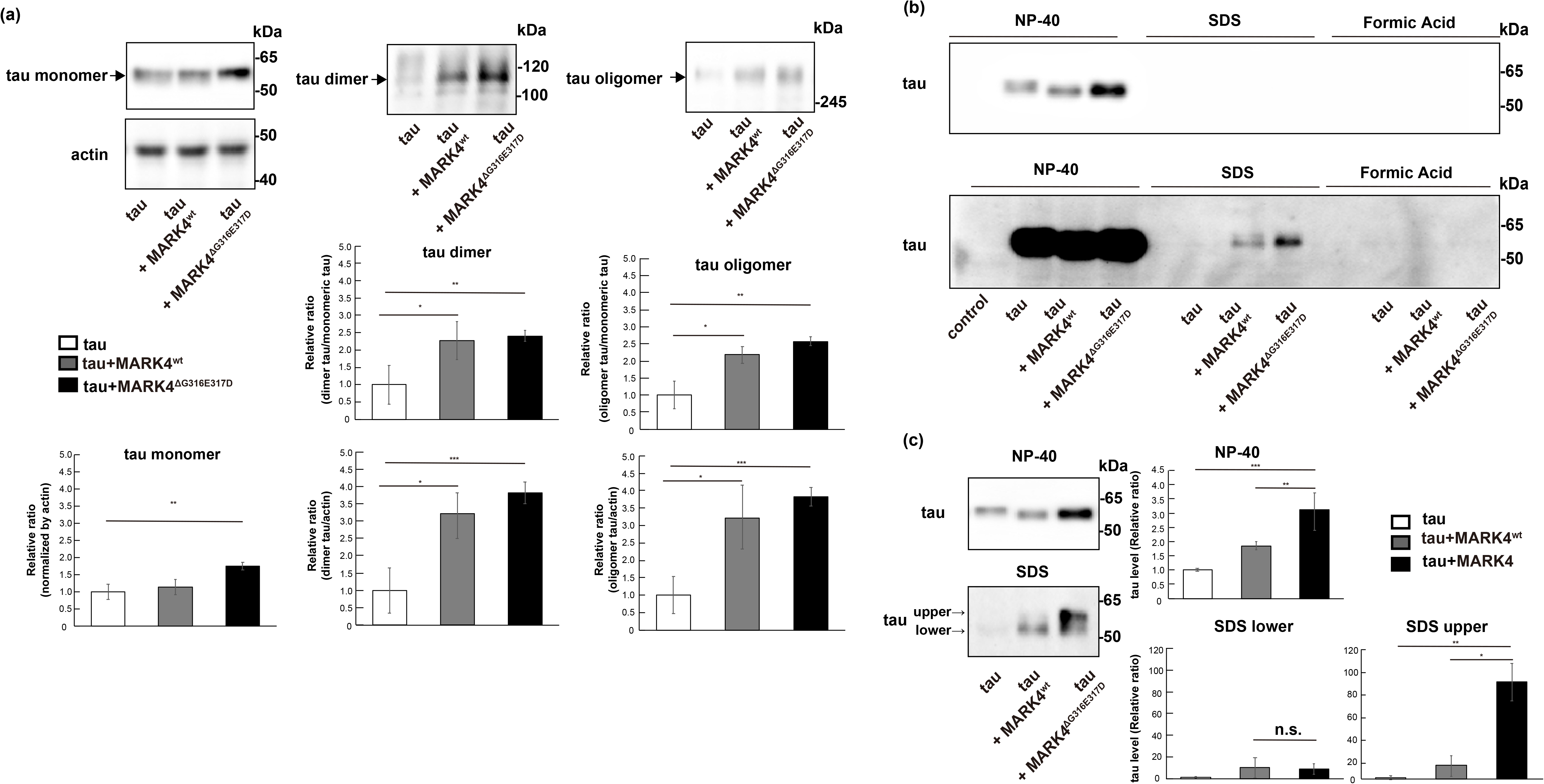
MARK4^ΔG316E317D^ promotes misfolding of tau. (a) MARK4^wt^ and MARK4^ΔG316E317D^ promoted accumulation of tau oligomers and dimers at similar levels. Western blotting was performed on fly head lysates prepared under non-reducing conditions with oligomer-specific antibody (oligomer) and pan-tau antibody (dimer). Representative blots and quantification are shown. Actin was used as a loading control. Means ± SEM; n = 3. n.s., p > 0.05; * p < 0.05, ** p < 0.01 (one-way ANOVA and Tukey post-hoc test). (b, c) Tau in extracts from heads expressing tau alone (tau), co-expressing tau and MARK4^wt^ (tau+MARK4^wt^), or co-expressing tau and MARK4^ΔG316E317D^ (tau+MARK4^ΔG316E317D^) were detected by western blot. Head homogenates were extracted first with RIPA buffer containing NP-40 (NP-40), next with 2%SDS (SDS), and finally with 70% formic acid (FA). (b) Representative western blots are shown (shorter exposure (upper panel) and longer exposure (lower panel)). (c) Western blotting was performed on fly heads extracted with RIPA buffer containing NP-40 (NP-40) and then with 2% SDS (SDS). Representative blots (left) and quantification (right) are shown.

Next, to assess whether co-expression of MARK4^wt^ or MARK4^ΔG316E317D^ would affect the detergent insolubility of tau, we performed sequential extraction followed by western blotting with anti-tau antibody (35). As reported previously, tau proteins in *Drosophila* retina exist mostly in detergent-soluble form ((19, 34), Figure 7b). When expressed alone or co-expressed with MARK4^wt^ or MARK4^ΔG316E317D^, tau was mostly extracted with RIPA buffer containing NP-40 (Figure 7b, NP-40). However, tau proteins were present in the SDS-soluble fraction when tau was co-expressed with MARK4^wt^ or MARK4^ΔG316E317D^, and this effect was more prominent when MARK4^ΔG316E317D^ was expressed (Figure 7b, SDS). Tau was not detected in the detergent-insoluble fraction when co-expressed with either MARK4^wt^ or MARK4^ΔG316E317D^ (Figure 7b, FA). Tau species in the NP-40 fraction were mostly tau_lower_ (Figure 7c, NP-40), whereas tau proteins accumulating in the SDS fraction contained both tau^upper^ and tau_lower_. Consistent with that result shown in Figure 6b, co-expression of MARK4^ΔG316E317D^, but not MARK4^wt^, increased the level of tau^upper^ in the SDS-soluble fraction (Figure 7c, SDS).

Taken together, these results indicate that MARK4^wt^ increases the levels of Ser262/356-phosphorylated tau, dimers, and oligomers. By contrast, MARK4^ΔG316E317D^ promotes accumulation of not only those tau species, but also of highly phosphorylated and insoluble tau species, which may promote neurodegeneration.

## Discussion

Accumulating evidence suggests that Par-1/MARK plays an initiating role in tau abnormalities leading to disease pathogenesis (13–19). We previously reported that Par-1 overexpression increases, and knockdown decreases, tau phosphorylation at Ser262 and Ser356, which leads to accumulation of tau and exacerbation of neurodegeneration in a fly model of tau toxicity (13, 14). In this study, we found that MARK4^wt^ increased the levels of pTauSer262, pTauSer356, and total tau, as well as increased tau toxicity (Figure 1 and 2). However, MARK4^wt^ did not either increase S2A tau levels or affect its toxicity (Figure 4), suggesting that MARK4^wt^ affects tau metabolism through tau phosphorylation at Ser262 and Ser356. On the other hand, we found that MARK4^ΔG316E317D^ promoted tau accumulation via additional mechanisms. MARK4^ΔG316E317D^ increased S2A tau levels (Figure 4), indicating that MARK4^ΔG316E317D^ can increase the abundance of tau that is not phosphorylated at Ser262 or Ser356. We also found that MARK4^ΔG316E317D^, but not MARK4^wt^, caused accumulation of the highly phosphorylated, aggregated forms of tau (Figure 6 and 7). These results suggest that MARK4^wt^ promotes accumulation of tau phosphorylated at Ser262 and 356, while MARK4^ΔG316E317D^ additionally increases accumulation of highly phosphorylated and more misfolded tau species.

Rovelet-Lecrux et al. reported that when HEK293 cells were co-transfected with tau and either MARK4^wt^or MARK4^ΔG316E317D^ constructs, cells transfected with MARK4^ΔG316E317D^ exhibited a significant increase in tau phosphorylation at Ser262 relative to MARK4^wt^. The authors of that study concluded that this *de novo* mutation in MARK4 results in a gain-of-function in the ability of MARK4 to phosphorylate tau on Ser262 (23). We also found that co-expression of MARK4^ΔG316E317D^ increased tau phosphorylated at Ser262 to a greater extent than MARK4^wt^ (Figure 1). However, *in vitro* kinase assays revealed that MARK4^wt^ and MARK4^ΔG316E317D^ did not significantly differ in terms of kinase activity (Figure 3), arguing against the idea that accumulation of tau is the result of more efficient phosphorylation of tau by MARK4^ΔG316E317D^.

The double mutation (ΔG316E317InsD) is located in the short linker that tethers the kinase to substrates or regulators (23). The predicted three-dimensional structure suggests that the double amino-acid change (ΔG316E317InsD) increases the area of the substrate-binding region, facilitating a strong association of MARK4 with interacting proteins (23). The strengthened interaction of MARK4^ΔG316E317D^ and substrates other than tau, or interaction with novel substrates, may indirectly stabilize highly phosphorylated misfolded forms of tau.

In addition to AD, MARK4 has been suggested to play a critical role in other neurodegenerative diseases, including ischemic axonal injury (21) and synucleinopathies (36). Under pathological conditions, abnormalities in MARK4, such as additional post-translational modifications in the linker region, might cause effects similar to those of the ΔG316E317InsD mutation on MARK4, and thus promote misfolding of tau and other aggregation-prone proteins. Identification of ΔG316E317D-induced changes in MARK4^ΔG316E317D^ that promote tau accumulation may provide insight into the dysregulation of MARK4 in the pathogenesis of sporadic AD.

## Materials and Methods

### Fly stocks and husbandry

Flies were maintained in standard cornmeal media at 25°C under light-dark cycles of 12:12 hours. The transgenic fly line carrying the human 0N4R tau, which has four tubulin-binding domains (R) at the C-terminal region and no N-terminal insert (N), is a kind gift from Dr. M. B. Feany (Harvard Medical School) (37). UAS-PAR1 RNAi is a kind gift from Dr. Bingwei Lu (Stanford University) (15). UAS-S2A tau were reported previously (13, 14, 18). To establish transgenic fly lines carrying UAS-MARK4^wt^ and UAS-MARK4^ΔG316E317D^, cDNA coding human MARK4 was obtained from DNASU Plasmid Repository (Clone# HsCD00294885, The Biodesign Institute at Arizona State University). ΔG316E317D mutation was introduced by using PCR-based site-directed mutagenesis. cDNA were subcloned into pUASTattB, and injected to the oocytes carrying into P{CaryP}attP2. Genotypes of the flies used in the experiments are described in Table S1.

### Chemicals and antibodies

An anti-MYC antibody (4A6) (Merck Millipore), anti-tau (T46) (Thermo), anti-tau (Tau5) (Thermo Fisher), anti-phospho-Ser262 tau antibody (Abcam), anti-phospho-Thr231 tau antibody (AT180, Thermo Fisher), anti-phospho-Ser356 tau antibody (Abcam), anti-oligomer antibody (T22) (Merck Millipore), anti-phospho-S6K (Cell Signaling), anti-Ref2p (Abcam), anti-LC3 (Merck Millipore), and anti-actin antibody (SIGMA) were purchased. AT8 antibody and PHF1 antibody are kind gift from Dr. Peter Davis (Albert Einstein College of Medicine).

### Cell culture and transfection

HEK293 cells were maintained in Dulbecco’s modified Eagle’s medium (DMEM, Sigma) supplemented with 10% (v/v) fetal bovine serum, 100 U/ml penicillin, and 0.1 mg/ml streptomycin. Plasmids encoding MARK4 was transfected with Lipofectamine 2000 (Thermo Fisher) according to the manufacturer’s protocol.

### In vitro kinase assay of MARK4

MARK4 expressed in HEK293 cells was immunoprecipitated from the cell lysate with monoclonal anti-Myc antibody (4A6) and Dynabeads protein G (Thermo Fisher). Its kinase activity was measured using human 2N4R tau, which was expressed and purified from E. coli and [γ32P]ATP as substrates. Incorporation of 32P into tau was quantified using liquid scintillation counter (Beckman).

### SDS-PAGE, Phos-tag SDS-PAGE, and immunoblotting

SDS-PAGE for western blotting of tau, MARK4 and actin was performed using 10% or 7.5% (w/v) polyacrylamide gels. Phos-tag SDS-PAGE was performed using 7.5% (w/v) polyacrylamide gels containing 25 μM or 50 μM Phos-tag acrylamide (Wako Chemicals) for tau, Proteins separated in the gel were transferred to a PVDF membrane (Merck Millipore) using a submerged blotting apparatus and then visualized using Immobilon Western Chemiluminescent HRP Substrate (Millipore). The chemiluminescent signal was detected by Fusion FX (Vilber) and intensity was quantified using ImageJ (NIH). Western blots were repeated a minimum of three times with different animals and representative blots are shown. Flies used for Western blotting were 2 day-old after eclosion.

### Histological analysis

Neurodegeneration in the fly retina was analyzed as previously reported (30). Fly heads were fixed in Bouin’s fixative for 48 hr at room temperature, incubated for 24 hr in 50 mM Tris/150 mM NaCl, and embedded in paraffin. Serial sections (7 μm thickness) through the entire heads were stained with hematoxylin and eosin and examined by bright-field microscopy. Images of the sections that include the lamina were captured with Keyence microscope BZ-X700 (Keyence), and vacuole area was measured using ImageJ (NIH). Heads from more than three flies (more than five hemispheres) were analyzed for each genotype

### qRT-PCR

qRT-PCR was carried out as previously reported (30). More than thirty flies for each genotype were collected and frozen. Heads were mechanically isolated, and total RNA was extracted using Isogen Reagent (NipponGene) according to the manufacturer’s protocol with an additional centrifugation step (11,000 x g for 5 min) to remove cuticle membranes prior to the addition of chloroform. Total RNA was reverse-transcribed using PrimeScript Master Mix (Takara Bio). qRT-PCR was performed using TOYOBO THUNDERBIRD SYBR qPCR Mix on a Thermal Cycler Dice Real Time System (Takara Bio). The average threshold cycle value (CT) was calculated from at least three replicates per sample. Expression of genes of interest was standardized relative to rp49. Relative expression values were determined by the ΔΔCT method. Primers were designed using Primer-Blast (NIH). The following primers were used for RT-qPCR:

htau for 5′-CAAGACAGACCACGGGGCGG-3′
htau rev 5′-CTGCTTGGCCAGGGAGGCAG-3′
rp49 for 5′-GCTAAGCTGTCGCACAAATG-3′
rp49 rev 5′-GTTCGATCCGTAACCGATGT-3′

### Sequential extraction of tau protein

The following sequential series of solubilizing buffer are RIPA (50 mM Tris-HCl, pH 8.0, 150 mM NaCl, 20 mM EDTA, 1& NP-40, and 50 mM NaF), 2%SDS, and 70% Formic acid (FA) (35). The sample were mixed with the indicated buffer, sonicated briefly, and centrifuged at 45,000 x g for 30 min at 4 degrees. The pellet was subjected to extraction with the indicated buffer. The supernatants were collected and subjected to immunoblot analysis as described above. The same volume of each extract was loaded per lane.

### Statistics

Statistics were done with Microsoft Excel (Microsoft) and R (R Foundation for Statistical Computing, Vienna, Austria.URL http://www.R-project.org/). Differences were assessed using the Student’s t-test or One-way ANOVA and Tukey’s honestly significant difference (HSD) post hoc test. P values < 0.05 were considered statistically significant.

## Competing interests

Authors declare no competing interests.

## Acknowledgement

The authors thank Drs. Mel Feany and Bingwei Lu; TRiP at Harvard Medical School (NIH/NIGMS R01-GM084947); the Bloomington Stock Center; and NIG Drosophila stock center, for fly stocks. We thank Dr. Peter Davis for AT8 and PHF1 antibody. We thank T. Miyashita for technical assistance. We thank Dr. S.-I. Hisanaga for critical comments. This work was supported by a Grant-in-Aid for Scientific Research on Innovative Areas (Brain Protein Aging and Dementia Control) [JSPS KAKENHI Grant number 17H05703] (to K.A.), a research award from the Hoan-sha Foundation (to K.A.), the Takeda Science Foundation (to K.A.), a research award from the Japan Foundation for Aging and Health (to K.A) and a Grant-in-Aid for Scientific Research on Challenging Research (Exploratory) [JSPS KAKENHI Grant number 19K21593] (to K.A.), and the Research Funding for Longevity Science 19-7 from the National Center for Geriatrics and Gerontology, Japan (to K.M.I.).

